# AligNet: Alignment of Protein-Protein Interaction Networks

**DOI:** 10.1101/551242

**Authors:** R. Alberich, A. Alcalá, M. Llabrés, F. Rosselló, G. Valiente

## Abstract

One of the most difficult problems difficult problem in systems biology is to discover protein-protein interactions as well as their associated functions. The analysis and alignment of protein-protein interaction networks (PPIN), which are the standard model to describe protein-protein interactions, has become a key ingredient to obtain functional orthologs as well as evolutionary conserved pathways and protein complexes. Several methods have been proposed to solve the PPIN alignment problem, aimed to match conserved subnetworks or functionally related proteins. However, the right balance between considering network topology and biological information is one of the most difficult and key points in any PPIN alignment algorithm which, unfortunately, remains unsolved. Therefore, in this work, we propose AligNet, a new method and software tool for the pairwise global alignment of PPIN that produces biologically meaningful alignments and more efficient computations than state-of-the-art methods and tools, by achieving a good balance between structural matching and protein function conservation as well as reasonable running times.

## Background

The regulation of cellular processes is one of the most enigmatic topics in cell biology [18]. The activity of cellular life relies on the proper functioning of the extremely complex networks of interactions among numerous intracellular constituents. It is clear that proteins are the most active participants in those processes [15], and this is the reason why researchers invest so much effort in the study of proteins, from their shape classification to their interactions networks, trying to unveil their function.

The increasing amount of available data in almost all biological research areas and, in particular, on protein structure and protein-protein interaction networks (PPIN), encourages data analysis as an important tool to derive biological meaning. To understand the mechanisms of protein-protein recognition at the molecular level and to unravel the global picture of their interactions in the cell, many experimental techniques have been developed so far. Some methods characterize individual protein interactions, while others screen for interactions at a genome-wide scale. Some good recent surveys on this topic are [32, 35, 17].

In order to analyze and contrast the available data on PPIN, several pairwise alignment algorithms have been defined in the last 15 years. The early alignment algorithms detected similarities between small subnetworks [16, 20, 22, 23, 26]. PathBLAST [16] is a tool developed to search for specific pathways in a PPIN. In contrast to PathBLAST, NetAlign [23] is a web-based tool designed to identify the conserved network substructures between two PPIN. The method used in this tool and in the algorithm presented in [26] is the matching of isomorphic subgraphs. The pairwise alignment method introduced in [20], MaWISh, produces a local alignment of two PPIN by evaluating the similarity of their graph structures through a scoring function that accounts for evolutionary events, while the algorithm introduced in [22] to align PPIN is based on both protein sequence similarity and network topology similarity, and it uses integer quadratic programming. In addition, a new and efficient approach to obtain multiple local alignments is described in [10], based on conserved functional modules. All these methods and algorithms are able to detect and obtain similarities between subnetworks. Actually, the aim of local alignment algorithms is to find regions with the same network structure in the networks under comparison. For every region in one network, an alignment with some region in the other network may be obtained, but it may happen that these local alignments are mutually inconsistent, because the same protein in one network may be matched by different local alignments to different proteins in the other network: then, as a final result, it may happen that these local alignments cannot be extended to a global alignment of the pair of PPIN.

In contrast to these local alignment methods, a global alignment algorithm is aimed at finding the best overall alignment between whole PPIN [9]. A global alignment is then a matching mapping between the sets of proteins of two PPIN, in such a way that each protein in one network is matched to one, and only one, protein in the other network. The motivation to perform a global network alignment is to compare interactomes, and to understand cross-species variations [12]. Global network alignment is also related to the detection of functional orthologs [36]. The identification of orthologous groups is useful for genome annotation, studies on gene/protein evolution, comparative genomics, and the identification of taxonomically restricted sequences. Nevertheless, there are often proteins in a network that have no biologically meaningful correspondence in another network and, thus, a meaningful alignment between a pair of networks should not necessarily cover all of them.

The first algorithm for the global alignment of PPIN was IsoRANK [36]. This algorithm produces a matching between a pair of input networks, based on the idea that a protein in one network should be matched to a protein in the other network if, and only if, the neighbors of the two proteins can also be matched. In order to obtain the matching, the algorithm associates a score with each possible pair of nodes of the two networks, capturing the similarity of their neighborhoods. Then, the highest scoring matching is obtained. Thus, the final result of the IsoRANK algorithm is a global alignment of two networks, but the idea behind this algorithm is that two networks are similar if the network topologies are similar. However, since nodes in these networks correspond to proteins, the protein similarity should also be taken into account when matching two proteins; for instance, if two proteins share a similar sequence and have a similar topology in the corresponding networks, then their matching probability should be higher than for those with very different sequences.

The right balance between network topology and biological information is one of the most difficult and key points in any PPIN alignment algorithm. In fact, IsoRANK considers the possibility of taking biological information of the nodes (proteins) into account, by tuning a parameter in the score values. Several other algorithms for the global alignment of PPIN have been proposed based on the idea that “two nodes are similar if their corresponding neighbors are so,” and hence considering mainly network topology but also some biological features [23, 1, 27, 29, 14]. As a result, some of them obtain a high number of conserved interactions but a very low functional consistence between the matched proteins [27, 29]. Indeed, as it is stated in [8], after performing a comparison of existing algorithms for the pairwise alignment of PPIN, the evaluated algorithms have dramatic differences in the quality of the alignments they produce, either yielding good topological or good biological matchings, but few of them do well in both aspects. In addition, they are not efficient from the computational point of view: for some of them, the software system is not well organized and they can run out of memory or spend a lot of time. Moreover, even if they do produce alignments, tend to be meaningless since the coincidences among them are very poor.

Consider also the analysis of eight recent aligners (NATALIE [19], SPINAL [2], PISwap [6], MAGNA[37], HubAlign [14], L-GRAAL [25], OPTNET[7], and ModuleAlign[13]) performed in [24], where several topological scores (node coverage, topological coherence, induced conserved substructure and symmetric sub-structure) and different biological coherence scores (KEGG pathway annotations, Gene Ontology annotation) are considered. The authors of this study conclude that the agreement between the alignments produced by any two different aligners is very low (around 20%) and also that the topological scores are not in agreement with the biological coherence of the alignments. Even more, when the alignment process is guided by topological information only, they produce alignments with the highest topological coherence but the lowest biological coherence. In contrast, when alignments are guided by sequence information only, they produce alignments with the highest biological coherence but the lowest topological coherence. This becomes extremely inconvenient in those aligners where the user has to choose the value of a parameter in order to specify the desired balance between the topological and the sequence similarities.

Therefore, the election of the right alignment tool depends on the purpose of the alignment itself. If the aim of the alignment is to infer biological information about relationships between the proteins in the networks, then the aligner with highest functional coherence score must be considered. If the user is interested in finding conserved network substructures, then the aligner with highest topological score must be considered. However, it should be noticed that when we try to obtain a global alignment between two topologically different networks, like for instance a dense network with a high number of interactions and a sparse network with a low number of interactions, the maximum expected value of any topological score is very low. Thus, the topological score as a measure of alignment correctness should be considered mainly to detect small conserved subnetworks, while, instead, the functional coherence score should be considered as the best measure of correctness in a global network alignment, whose goal is the detection of functional orthologs.

Motivated by the lack of well-balanced and efficient algorithms, we have designed AligNet, a parameter-free PPIN alignment algorithm aimed at filling the gap between efficient topologically and biologically meaningful matchings. The overall idea of the algorithm is to obtain many local alignments that are combined and extended into a meaningful global alignment. The final alignment captures the benefits of considering both categories of alignments. With the local alignments we capture the topological similarity between the networks and we speed up the running time of the algorithm, while with the final global alignment we solve the inconsistencies among the local alignments and yield an overall alignment of the pair of input PPIN. The results obtained with AligNet and with the best aligners compared in [8] and in [24] show that AligNet indeed achieves a good balance between topological and biological matching. In the tests reported in this paper, AligNet obtained the highest functional coherence scores in most of the alignments, which means that it maximized the functional consistence between aligned proteins, and also a reasonable fraction of conserved interactions. In addition, HubAlign and AligNet had the best running times among all the aligners considered in the aforementioned tests.

## Methods

### Protein-protein interaction networks as graphs

A protein-protein interaction network (PPIN) is modelled in a natural way as a graph, with its nodes representing the network’s proteins and its edges, the interactions between them. Moreover, the interaction between two proteins is considered a symmetric property, that is, if a protein *p*_1_ interacts with another protein *p*_2_, then it is tacitly understood that *p*_2_ also interacts with *p*_1_. Hence, PPIN are specifically modelled by means of undirected graphs. In this way, the problem of aligning pairs of PPIN is translated into the problem of aligning pairs of undirected graphs with their nodes injectively labelled by proteins.

Formally, an (*undirected*) *graph* is a structure *G* = (*V*, *E*) with *V* a finite set of *nodes* and *E* a family of 2-element subsets {*u*, *v*} of *V*, called the *edges* of the graph; recall that, as sets, {*u*, *v*} = {*v*, *u*}. We say that an edge *e* = {*u*, *v*} *connects* the nodes *u* and *v*, and also that *e* is *incident* to *u* and *v*. The nodes *v* such that {*u*, *v*} ∈ *E* are the *neighbors* of *u*. We shall denote by *N_G_*(*u*) the set of neighbors of *u* in *G*.

We introduce now some further definitions and notation that will be used throughout this paper. Let *G* = (*V*, *E*) be an undirected graph.

- The *degree* of a node *u* ∈ *V* is the number of edges incident to it; that is, the cardinal of *N_G_*(*u*). We denote it by deg(*u*).
- A *path* between two nodes *u*, *v* ∈ *V* is a sequence of pairwise different edges {*u*, *u*_1_}, {*u*_1_, *u*_2_}, …, {*u*_*k*−1_, *u_k_*}, {*u_k_*, *v*} such that the first and last edges are incident to *u* and *v*, respectively, and every pair of consecutive edges share a node (different from *u* and *v*, in the case of the first and last edges, respectively). Two nodes are *connected* when there exists a path between them. The *length* of a path is the number of edges forming it, and its *intermediate nodes* are *u*_1_, …, *u_k_*.
- For every pair of connected nodes *u*, *v* ∈ *V*, their *distance* in *G* is the length of a shortest path connecting them. We denote it by *d_G_*(*u*, *v*).
- The *diameter* of *G* is the maximum distance between any two connected nodes in *G*. We denote it by *D*(*G*).

Figure 1 displays two toy PPIN that will be used as a running example throughout this section. The first network consists of 8 nodes and 9 edges, while the second network consists of 9 nodes and 17 edges.

**Figure 1:**
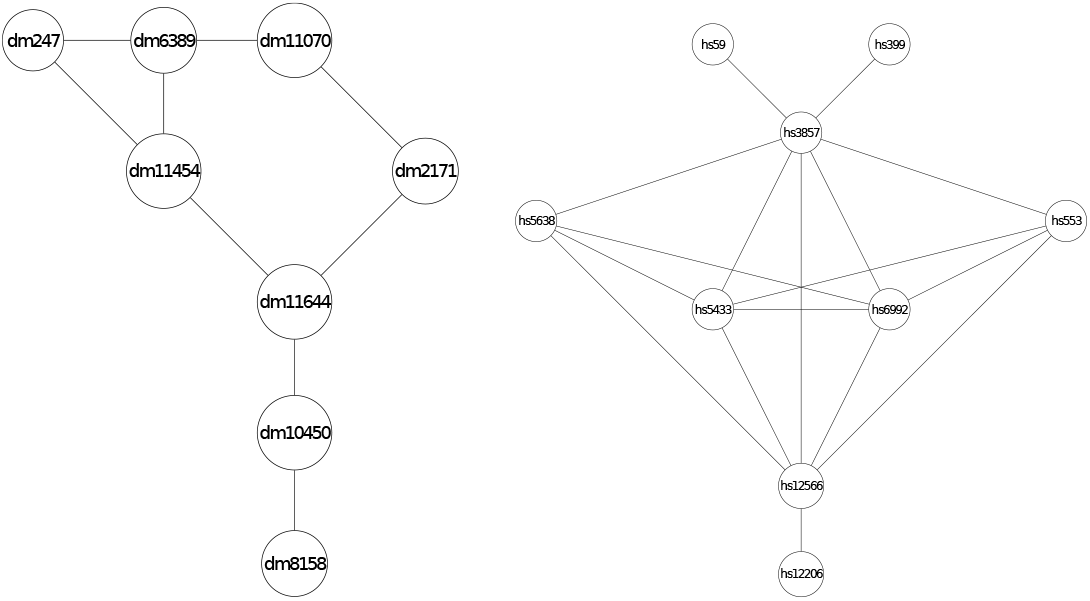
This figure shows two small pieces of PPINs that we shall use to visualize the performance of AligNet. The subnetworks belong to the *Drosophila melanogaster* (dme) and the *Homo sapiens* (hsa) PPINs contained in the IsoBase database. The first network has 8 proteins and 9 interactions, and the second network has 9 proteins and 17 interactions. The diameter of the first network is 4 and the diameter of the second network is 3.

### The structure of the AligNet algorithm

AligNet receives as input two graphs *G* and *G*′ representing two PPIN (in particular, each node of them is labeled with a protein, in such a way that different nodes in a graph correspond to different proteins) and produces, as output, a similarity score for them and a local and a global alignment between them. AligNet has been implemented in R [31], and the implementation is freely available from http://bioinfo.uib.es/~recerca/AligNet/.

The main steps in AligNet are:

1. The computation of overlapping clusterings *C*(G) and *C*(*G*′), respectively, of the input networks *G* and *G*′.
2. The computation of alignments between pairs of clusters in *C*(*G*) and *C*(*G*′).
3. The computation of a matching between *C*(*G*) and *C*(*G*′).
4. The computation of a local alignment of the input networks *G* and *G*′.
5. The extension of this local alignment to a meaningful global alignment.

Throughout this section, *G* = (*V*, *E*) and *G*′ = (*V*′, *E*′) will denote two graphs representing the input PPIN. We shall identify each node in any of these graphs with the protein it represents.

#### Step 1. Overlapping clusterings

The first step in AligNet consists in computing an overlapping clustering of each input network. These clusterings are based on a specific similarity score *s*(*u*, *v*) between pairs of proteins (nodes) *u*, *v* in a PPIN, which is defined as follows: for every pair of connected nodes *u*, *v* in a graph G representing a PPIN,

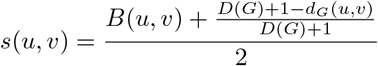

where:

- *D*(*G*) is the diameter of the *G* and *d_G_*(*u*, *v*) is the distance between *u* and *v*.
- *B*(*u*, *v*) is the *normalized bit score* of the proteins associated to the nodes *u* and *v*, that is, a rescaled version of their alignment score obtained with BLAST+, which is independent of the size of the search space [5].

If *u*, *v* are not connected by a path, then *s*(*u*, *v*) = 0.

The intuition behind this similarity score is that two proteins are similar if they have similar sequences of nucleotides and they are relatively close to each other in the graph. Recall that two proteins interact when there is an edge connecting them, that is, when their distance is 1. Therefore, the plausibility that two proteins have a related biological function increases when they are close to each other in the graph.

To obtain the overlapping clustering of an input network, we define a cluster centered at every node of the graph as follows. Let *α* be the third quartile of the distribution of the similarity score values of pairs of nodes: that is, *α* is the value for which only 25% of the pairs of nodes (*u*, *v*) are such that *s*(*u*, *v*) > *α*. Then, for every node *u* ∈ *V*, the *cluster C_u_* in *G centered* at *u* is

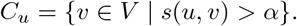

We denote by *C*(*G*) the set of clusters of a PPIN *G*.

So, the first step of AligNet computes the overlapping clusterings *C*(*G*) and *C*(*G*′) of the input networks *G* and *G*′. As a running example throughout this section we will consider two small networks. Figure 2 displays the first PPI network considered as a running example as well as its overlapping clustering. This first network consists of 8 nodes and 9 edges, so there are 8 clusters. Figure 2 displays the second PPI network which consists of 9 nodes and 17 edges, and its overlapping clustering has 9 clusters.

**Figure 2:**
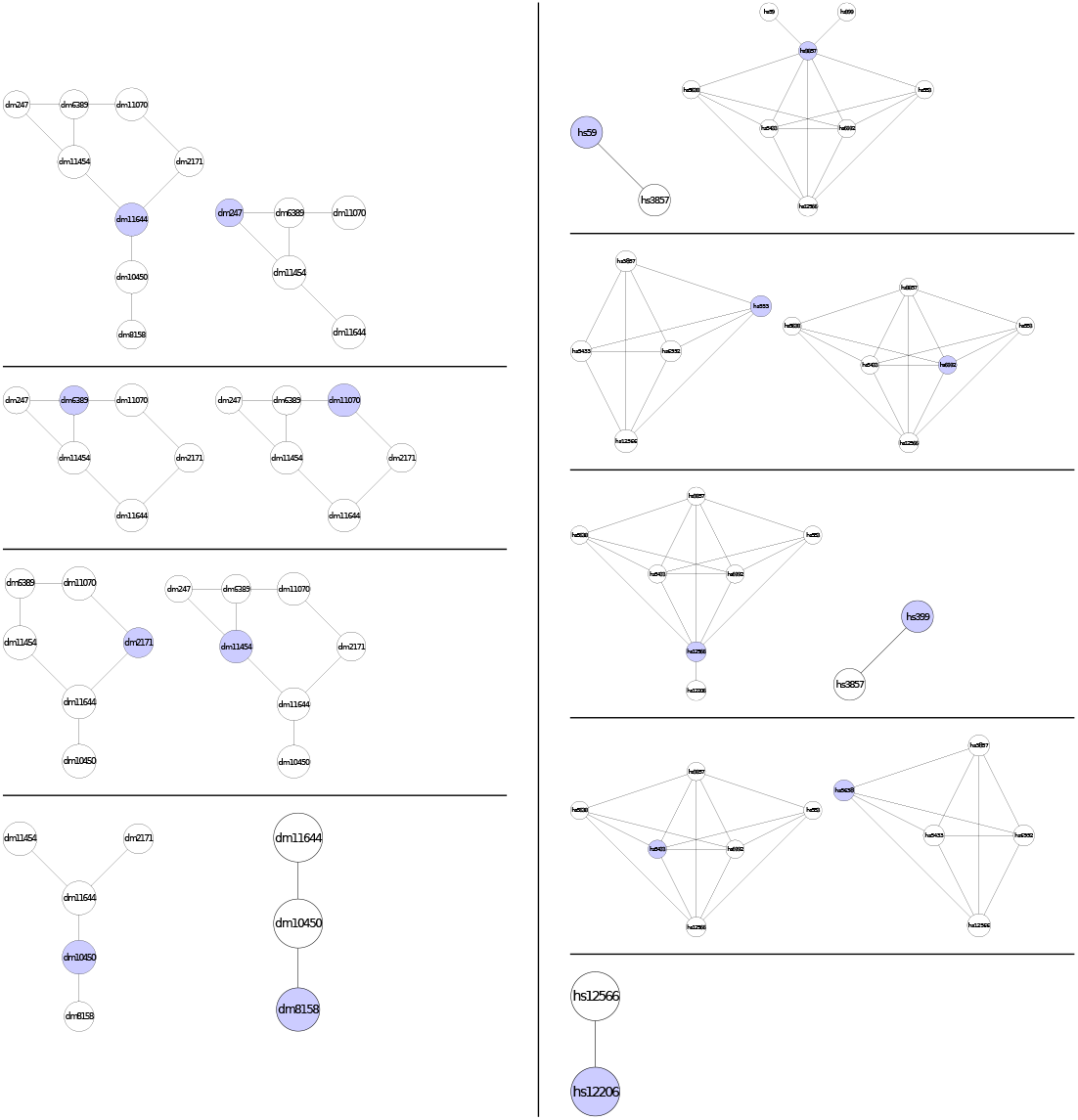
This figure shows the overlapping clustering on the PPINs in Figure 1 obtained by AligNet. We can see here the 8 clusters in the network in Figure 1 on the left, and the 9 clusters in the network in Figure 1 on the right. The center of every cluster is highlighted in blue. Since we have considered two small pieces of a PPIN, we obtain here that, the first cluster on the left is the entire piece of network. In the right, we obtain also the entire piece of network in the second cluster on the right. Notice that we obtain the whole piece of the network when we consider the cluster of a node that is in the center of the network..

#### Step 2. Alignments between pairs of clusters

In this second step, AligNet computes an alignment between every pair of clusters *C_u_* ∈ *C*(*G*) and *C*_*u*′_ ∈ *C*(*G*′) such that *B*(*u*,*u*′) > 0. That is, if we assume that *B*(*u*,*u*′) > 0 for every pair *u* ∈ *V* and *u*′ ∈ *V*′ then AligNet computes |*V*| · |*V*′| alignments. These alignments define an alignment score between every such a pair of clusters that will be used in the third step to compute a matching between *C*(*G*) and *C*(*G*′).

The general idea to obtain the alignment between a pair of clusters *C_u_* ∈ *C*(*G*) and *C_u′_* ∈ *C*(*G*′) (with *B*(*u*,*u*′) > 0) is the following: we first match the centers of the clusters, that is, we match *u* with *u*′ and then, we match the neighbors of *u* to the neighbors of *u*′. To decide the neighbors matching, we take into account their sequence similarity and their degrees. Thus, a neighbor of *u* is matched to a neighbor of *u*′ provided that they have similar nucleotide sequences and also similar degrees. Following the same criteria, we match the neighbors of the neighbors of *u* with the neighbors of the neighbors of *u*′. We iterate this process until no unmatched neighbors are found. In the intermediate steps we keep the node matching in a list of pairs denoted by *L*_*u*,*u*′_. When the algorithm terminates, *L*_*u*,*u*′_ provides a partial mapping between the nodes in *C_u_* and the nodes in 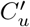.

Thus, the alignment between a pair of clusters *C_u_* ∈ *C*(*G*) and *C*_*u*′_ ∈ *C*(*G*′) (with *B*(*u*, *u*′) > 0) can be formally defined as follows:

i. Match *u* with *u*′. Set 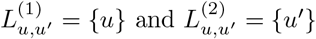.
ii. For every *v* ∈ *C_u_* ∩ *N_G_*(*u*) and for every *v*′ ∈ *C*_*u*′_ ∩ *N_G′_* (*u*′), let

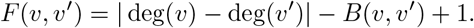 Compute a matching *M*_*u*,*u*′_ ⊆ (*C_u_* ∩ *N_G_*(*u*)) × (*C*_*u*′_ ∩ *N*_*G*′_(*u*′)) that minimizes ∑_(*v*, *v*′)∈*M*_*u*, *u*′__ *F*(*v*, *v*′). Sort the pairs in *M*_*u*, *u*′_ in decreasing order of their *F* value, and concatenate them to *L*_*u*,*u*′_. Add their first coordinates to 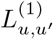 and their second coordinates to 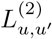.
iii. Iterate step (ii), replacing (*u*, *u*′) by the rest of the pairs in *L*_*u*, *u*′_ and removing from *C*_*u*_ and *C*_*u*′_ the nodes already aligned. More specifically, in the *k*-th iteration, take the *k*-th element 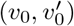 of *L*_*u*, *u*′_. For every 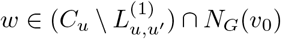 and every 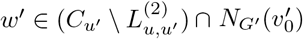, compute *F*(*w*, *w*′). Then, compute a matching

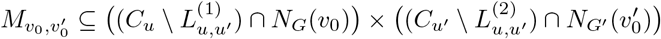

that minimizes 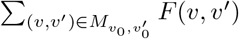. Sort the pairs forming 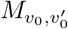 in decreasing order of their *F* value, and concatenate them to *L*_*u*, *u*′_. Add their first coordinates to 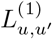, and their second coordinates to 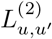.

The matchings in step (ii) as well as in each iteration in step (iii) are computed with the Hungarian algorithm [21]. Figure 3 shows an example of the alignment of a pair of clusters: one cluster from the first network and another cluster from the second network.

**Figure 3:**
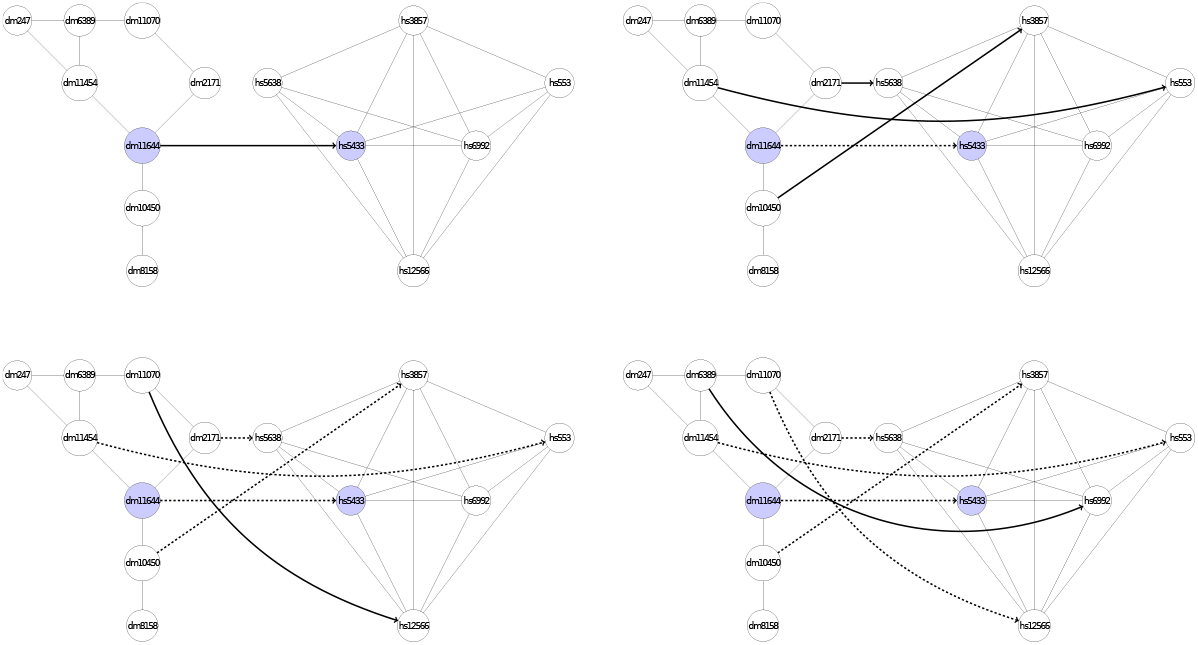
This figure shows how AligNet aligns two clusters which corresponds to Step 2 of our algorithm. The clusters in this example are, respectively, the first in the list of clusters of *G*, which are shown on the left in Figure 2 and the seventh in the list of clusters of *G*′, which are shown on the right in Figure 2. We show in the picture all the steps needed to align the cluster of *G* with the cluster of *G*′. From top to bottom in this figure, we can see that AligNet first aligns the centers of the clusters, which are the nodes highlighted in blue. Then, AligNet aligns the neighbors of the centers (second row). Next, AligNet aligns the neighbors of the neighbors. In each step we show in dashed lines the nodes that are already aligned and in solid lines the nodes that are aligned in the present step. Notice that, in this example, there are two nodes that remain unmatched.

The overall idea behind the algorithm described above is that a node *v* in *C_u_* should be matched to a node *v*′ in *C*_*u*′_ when they have a similar topological role in the cluster and similar sequences, provided that, furthermore, there exist paths connecting the cluster centers *u* and *u*′ with *v* and *v*′, respectively, such that their intermediate nodes are already aligned in sequential order along the paths. The alignment procedure gives priority to matching neighbors of nodes *x*, *x*′ at the possible shortest distance of the respective cluster centers and with *F*(*x*, *x*′) as large as possible among those pairs already matched at their same iterative step.

The resulting alignment *L*_*u*, *u*′_ defines a partial injective mapping *η*_*u*,*u*′_ : *C_u_* → *C*_*u*′_. The nodes in *C_u_* that are matched to nodes in *C*_*u*′_ form the domain of the mapping *η*_*u*,*u*′_, which is denoted by *Dom η*_*u*, *u*′_.

#### Step 3. Matching between families of clusters

Let

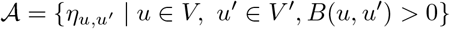

be the set of alignments obtained in step 2. The score of every alignment 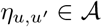 is defined as

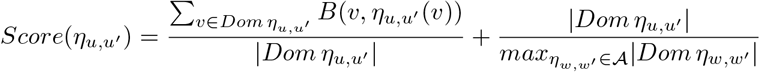

where |*X*| stands for the number of elements in the set *X*. This score assesses simultaneously the average similarity of the sequences of the proteins matched by *η*_*u*,*u*′_ and their number.

Once computed all these scores, AligNet obtains a matching between *C*(*G*) and *C*(*G*′) by considering a bipartite graph where the nodes are the clusters in *C*(*G*) and *C*(*G*′), the edges correspond to alignments 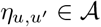, and the weight of an edge connecting *C_u_* with *C*_*u*′_ is the corresponding score *Score*(*η*_*u*, *u*′_). The matching between the nodes in *C*(*G*) and the nodes in *C*(*G*′) is then obtained by applying the maximum weighted bipartite matching algorithm to this bipartite graph. Recall that the nodes in *C*(*G*) are the clusters in *G* and the nodes in *C*(*G*′) are the clusters in *G*′. The solution to the maximum weighted bipartite matching problem provides us with a matching between the clusters in *G* and the clusters in *G*′. We shall denote by 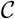 the set of partial injective mappings *η*_*u*,*u*′_ corresponding to pairs of clusters (*C_u_*, *C*_*u*′_) that are matched by this matching.

Figure 4 shows the matching between the family of clusters in Figure 1 and the family of clusters in Figure 2 obtained in this step.

**Figure 4:**
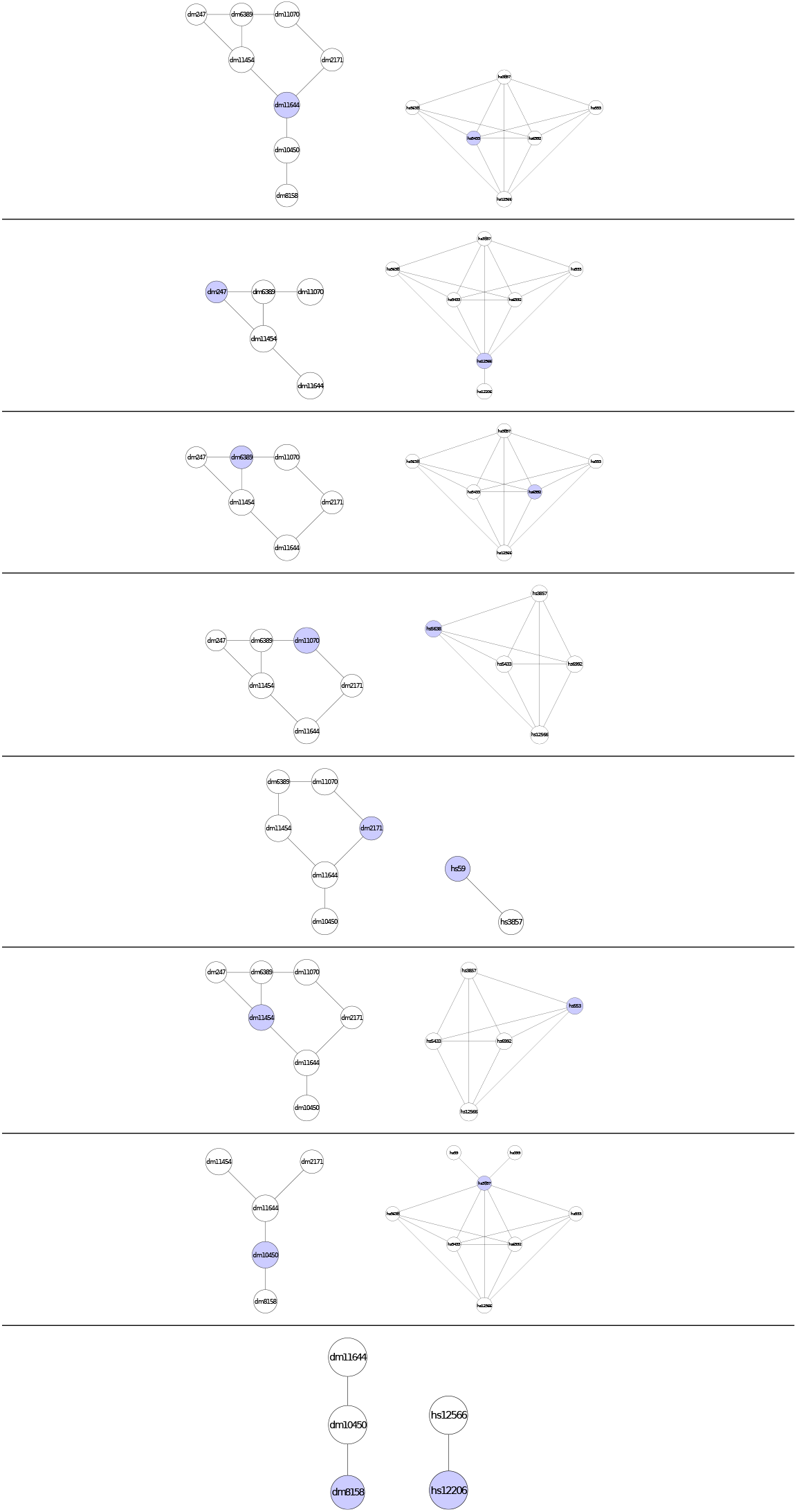
This figure shows the final assignment between the clusters in Figure 2 produced by AligNet, which corresponds also to Step 3. Each of the eight clusters obtained from *G* is aligned to one, and only one, of the nine clusters obtained from *G*′. Hence, one cluster from *G*′ remains unmatched which is the second cluster in the third row on the right in Figure 2. In this figure, we show the clusters from *G* on the left and its corresponding cluster image from *G*′ on the right.

#### Step 4. Local alignment of PPIN

In this step, AligNet produces a local alignment between *G* and *G*′ from the matching between *C*(*G*) and *C*(*G*′) obtained in the previous step. The main idea is to define this alignment by merging the partial injective mappings 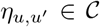. The problem is that these mappings may be inconsistent, because *C*(*G*) and *C*(*G*′) are overlapping clusterings. Indeed, it may happen that a node *w* belongs to more than one cluster *C_u_*, and that the corresponding mappings 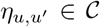 send *w* to different nodes in *G*′; conversely, for *w*′ belonging to multiple *C*_*u*′_ mappings to different nodes in *G*.

To overcome this problem, we consider a weighted bipartite hypergraph whose nodes are the nodes in *G* and in *G*′, every mapping *η*_*u*,*u*′_ is a hyperarc with source its domain and target its image, and the weight of every hyperarc is the score *Score*(*η*_*u*, *u*′_). Then, the solution of the weighted bipartite hypergraph assignment problem provides a well-defined local alignment of the input networks. However, in order to decrease the computation time of AligNet, we do not consider all the mappings *η*_*u*,*u*′_ together, but just a subset 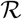 of them that is recursively increased until all mapping *η*_*u*,*u*′_ have been considered. Thus, AligNet builds recursively a subset 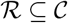 of *best-scored* alignments, by choosing, at each step, a mapping 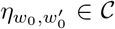 with *w*_0_ not belonging to the union of the domains of the mappings *η*_*w*, *w*′_ already in 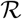 and with maximum 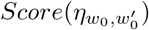 among all such mappings. AligNet iterates this procedure until every node in 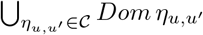 belongs to the domain of some mapping in 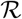. In Figure 5 we give the subset 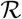 of 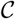 for the networks in our running example.

**Figure 5:**
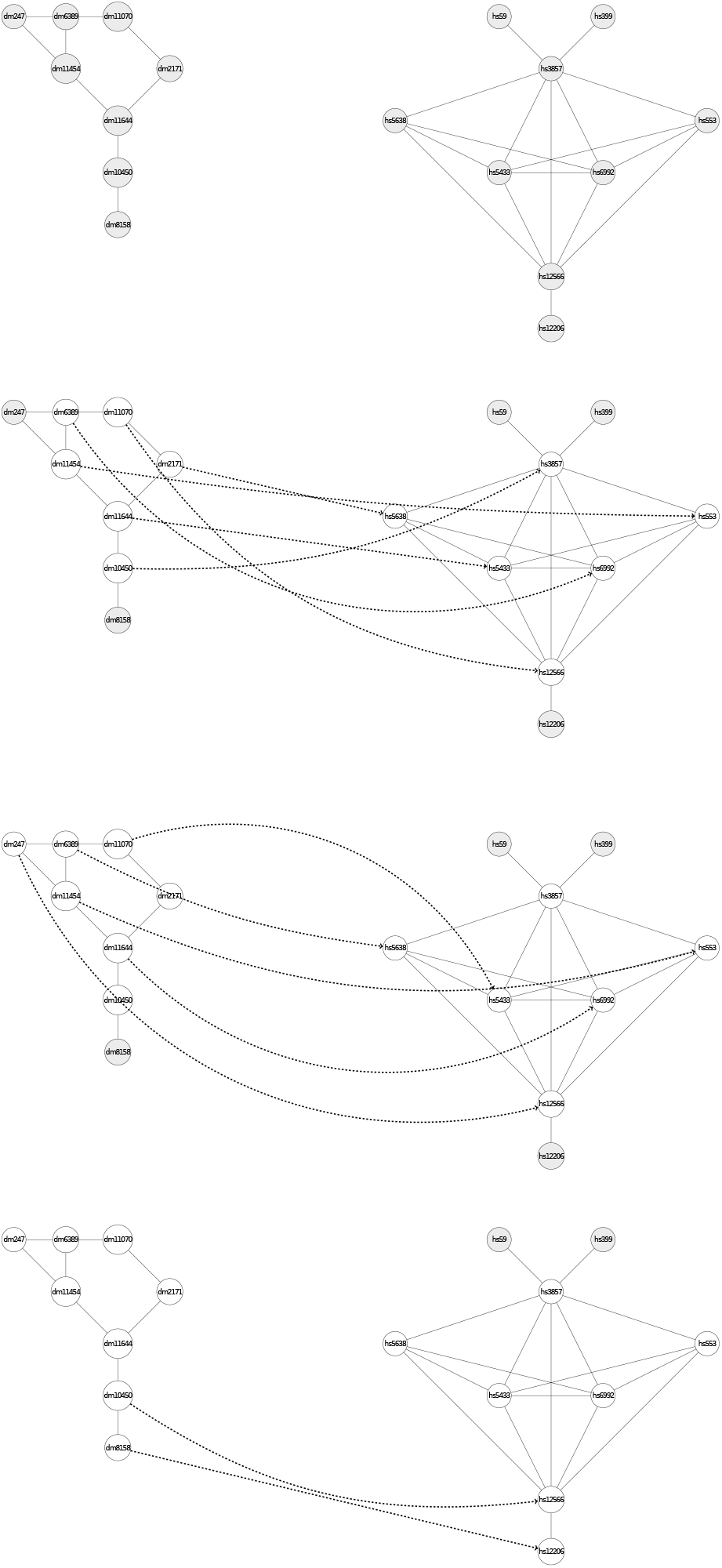
This figure shows how AligNet constructs an appropriate set of alignments considered to obtain a final local alignment. This corresponds to the Step 4 of our aligner. First of all, a maximum score alignment between a pair of clusters is chosen: in this case, this corresponds to the matching between the clusters in Figure 3. Both clusters are shown in the second row of this figure. The shadowed nodes are the nodes that are not aligned. Next, a maximum score alignment of a pair of clusters with source a cluster centered at a shadowed node is chosen: it turns out to be the one in the second row in Figure 4 and it is shown in the third row in this figure. Finally, the last alignment to be included in the appropriate set of alignments must be the one with source cluster centered at the remaining shadowed node: this corresponds to the alignment in the last row in Figure 4 shown in the bottom of this figure. Notice that in the end, that is when we consider the three alignments together, there are four nodes in the source network with inconsistent assignments..

Now, consider the directed hypergraph *H* with nodes *V* ∪ *V*′ and hyperarcs the mappings 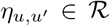: each *η*_*u*,*u*′_ is understood as a hyperarc with source its domain and target its image. Then, AligNet obtains from this hypergraph a local well-defined alignment between *G* and *G*′ as a solution of the corresponding weighted bipartite hypergraph assignment problem [4]. Figure 6 shows the local alignment obtained from the hypergraph corresponding to Figure 5.

**Figure 6:**
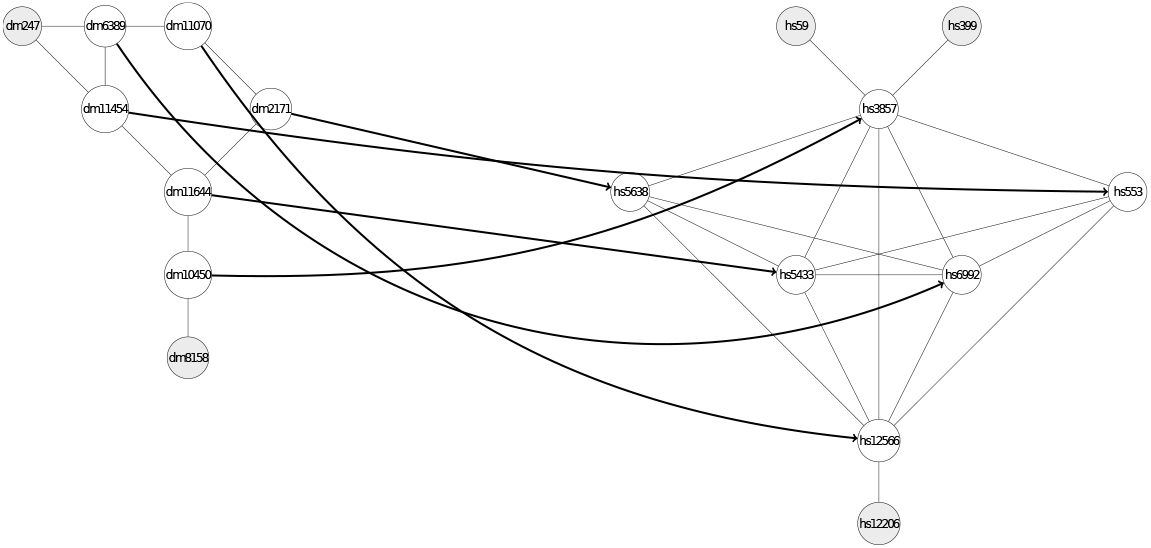
This figure shows the local alignment of the original networks obtained by AligNet in its Step 4, once the inconsistent assignments have been solved. The coherent assignment of nodes is obtained as the solution to the weighted bipartite hypergraph assignment problem, for the hypergraph associated to the appropriate set of alignments described in Figure 5. In this case, the hypergraph has three hyperarcs, corresponding to the three alignments considered in the appropriate set of alignments..

#### Step 5. Global meaningful alignment of PPIN

In order to extend the local alignment produced in the previous step, AligNet iterates the following procedure:

- It removes the nodes in *G* and *G*′ that have already been aligned, and it recomputes the score of each alignment *η*_*u*,*u*′_ following the same definition as in Step 3, but only taking into account the remaining nodes in its domain and image.
- It computes a new optimal matching 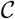 between *C*(*G*) and *C*(*G*′), as in step 3, but using as edges those *η*_*u*,*u*′_ whose updated score is positive, and weights these updated scores.
- It computes a new set 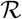 of best-scored alignments *η*_*u*,*u*′_ with *Score*(*η*_*u*, *u*′_) > 0, as in step 4.
- It defines a new directed hypergraph *H* whose nodes are the nodes in *V* ∪ *V*′ not yet aligned and hyperarcs the mappings *η*_*u*,*u*′_ in the new set 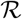, understood as hyperarcs with source the still unaligned nodes in their domain and target the still unaligned nodes in their image.
- It computes a local alignment between unaligned nodes in *V* and *V*′ by solving the weighted bipartite hypergraph assignment problem for this hypergraph, and it adds this local alignment to the alignment obtained so far.

This procedure is iterated while there exist nodes not aligned belonging to the domain or the image of some alignment *η*_*u*,*u*′_ with (updated) positive score: In Figure 7 we show the final global meaningful alignment obtained with AligNet for the networks in our running example.

**Figure 7:**
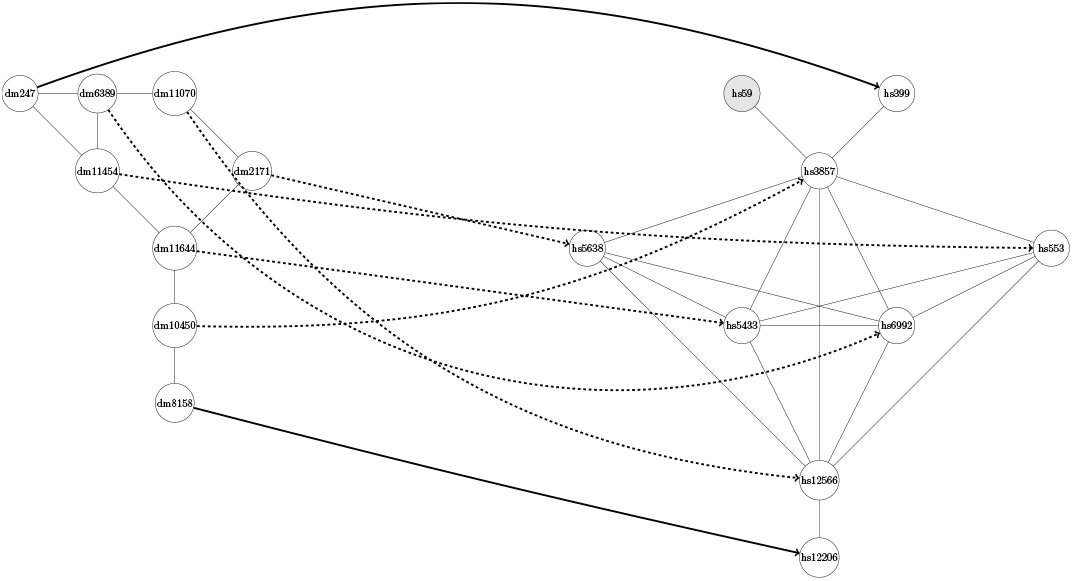
This figure shows the final global alignment of the original networks obtained by AligNet. Notice that, in Step 5 of AligNet, the previous alignment is extended to a global one. In this case, there were two unmatched nodes in the source network in Figure 6 which are now assigned. The assignment of these two nodes is shown with solid arrows while with dashed arrows we show the already assigned nodes.

### Evaluation of alignment quality

Several methods have been proposed to evaluate the quality of an alignment and to compare the performance of PPIN aligners [8, 24]. One of the handicaps when considering biological data is that the true alignment is unknown and, therefore, to evaluate the alignment quality one cannot count the number of true and false positives and true and false negatives. However, several measures of alignment quality have been already proposed which are divided in two categories, *topological coherence* and *biological coherence*. The topological coherence measures evaluate the topological similarity of the aligned regions considering the *edge correctness*, which is the percentage of conserved edges, the *induced conserved substructure*, which is the percentage of preserved edges, and the *symmetric substructure score*, which is the percentage of preserved and conserved edges. The biological coherence measures evaluate the protein function similarities of the aligned proteins by considering the *KEGG pathway annotation*, which measures the percentage of proteins aligned to proteins that participate in the same pathway, and the *Gene Ontology annotation*, which measures the similarity between the GO terms of a protein and its image. In the comparison of eight recent aligners reported in [24], it is stated that there is a strong correlation between the three topological coherence measures and also a strong correlation between the two biological coherence measures. Therefore, when evaluating the alignment quality, it is enough to consider one of the tree measures for topological coherence and one of the two biological coherence measures. However, there is a low correlation between the topological and the biological coherence measures. Thus, to evaluate the alignment quality, we considered both the edge correctness and the Gene Ontology annotation measures, which are defined as follows:

Let *G* = (*V*, *E*) and *G*′ = (*V*′,*E*′) be two PPIN such that |*V*| ⩽ |*V*′|. The *edge correctness ratio* of a mapping *μ* : *G* → *G*′ is the ratio of the edges that are preserved by *μ*, and it is defined by

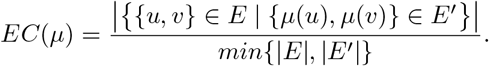

The *functional coherence value*, or *GO consistency*, of a mapping *μ* : *G* → *G*′ is defined as

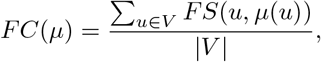

where the similarity score *FS* is defined by

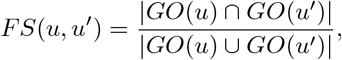

with *GO*(*u*) and *GO*(*u*′) the sets of GO annotations of the proteins *u* and *u*′, respectively.

## Results and Discussion

In this section we report the tests performed to assess the performance of AligNet. These tests have been designed taking into account the study and results reported in [8] and [24], where a comparison of algorithms for the pairwise alignment of biological networks, and in particular of PPIN, is given. Considering the conclusions reported in both studies, the aligners proposed among all the consider aligners because they produce better alignments are NATALIE [19], PINALOG [30], SPINAL [2], HubAlign [14] and L-GRAAL [25].

### Network data

We have considered the same dataset used in [8], so that it makes sense to compare the results obtained by AligNet with the results reported therein. Thus, we have downloaded from the IsoBase database [28] the PPIN (version 1.0.2) of five organisms: *M. musculus* (mus), *C. elegans* (cel), *D. melanogaster* (dme), *S. cerevisiae* (sce), and *H. sapiens* (hsa), and aligned each pair of them with AligNet. The number of nodes and edges of these PPIN are shown in Table 1.

**Table 1:** This table shows the number of nodes and edges, with and without loops, of the five networks considered as input data in our tests.

|  | Nodes | Edges (with loops) | E (without loops) |
| --- | --- | --- | --- |
| *M. musculus* | 623 | 776 | 559 |
| *C. elegans* | 2995 | 8639 | 4827 |
| *S. cerevisiae* | 5524 | 164718 | 82656 |
| *D. melanogaster* | 7396 | 49467 | 24937 |
| *H. sapiens* | 10403 | 105232 | 54654 |

### Comparison to other aligners

#### Quantitative analysis

In order to evaluate the quality of the alignments produced by AligNet, and to compare it with that of the aforementioned aligners, we have used both the edge correctness (EC) and the functional coherence (FC) metrics defined in the previous section.

As reported in [8], an edge correctness ratio of 100% should not be taken as a conclusive evidence of a correct network alignment, because it is always possible that two biologically unrelated edges had been mapped to each other. In addition, it is reported in [24] that the alignment of a dense network with a sparse network produces a low edge correctness ratio. We can observe that this situation is reflected for instance in the alignment between *S. cerevisiae* and *D. melanogaster*, since there are more edges in the source network (sce) than in the target network (dme). Thus, an edge correctness ratio of 20% should be considered an evidence of an incorrect network alignment, only when the number of edges of the source network is smaller than the number of edges in the target network. Therefore, given the limitations of this topological measure of alignment quality, measures of agreement derived from biological information are also popular in the literature. In particular, most papers on PPIN alignment make use of gene orthology annotations from the Gene Ontology (GO) database [3] to measure alignment accuracy, by comparing the similarity of GO annotations between aligned proteins. Hence, we have also used the functional coherence value, or GO consistency, introduced in the previous section, to assess the quality of our alignments.

In order to compare the EC scores and the FC scores of all the considered aligners for every pair of PPIN, we run all the aligners on the same pairs of input networks. However, as it has been already stated in previous studies of existing aligners [8, 24], some difficulties appear when trying to do this work. More precisely, there were computations that never stopped. This was the case of NATALIE. L-GRAAL matches the network with the smallest number of edges to the network with the largest number of edges, which means that it interchanges the order of the input networks in the case of the alignments between *S. cerevisiae* and *D. melanogaster* and between *S. cerevisiae* and *H. sapiens*. And also, for most of the aligners, some parameters must be fixed. We considered the parameters suggested by default in all the aligners whenever it was possible. With L-GRAAL, also a time limit or a maximum number of steps must be considered. We again decided to consider the parameters suggested by default in its implementation.

In Table 2 and Table 3 we report the results obtained with this test. We can observe there that the alignments of small networks with a low number of edges, such as *M. musculus*, produced alignments with high EC scores, especially when the target network has a large number of edges. However, even in this case, the EC scores obtained with the aligners PINALOG and SPINAL are not high. It is very surprising that SPINAL preserved less than 10% of the edges in the source network in all the alignments, except in the alignment between *M. musculus* and *H. sapiens*, only 24% of the edges present in *M. musculus* were preserved. On the other hand, PINALOG obtained much more reasonable scores than SPINAL but in the best scenario, that is, when the alignment is between *M. musculus* and the others networks, PINALOG preserved less than 40% of the edges in two alignments (*M. musculus* with *C. elegans* and *D. melanogaster*) while the other aligners preserved more than 60% of the edges present in *M. musculus*. The best scores in this case are obtained by HubAlign, which preserves more than 80% of the edges, followed by L-GRAAL that preserves the 70% of the edges, and then AligNet that preserves 60% of the edges. However, we can also observe here that, when the number of edges in the source network increases, the EC scores decrease dramatically even in the case of HubAlign. When considering the alignments between *S. cervisiae* and *D. melanogaster* or *H. sapiens*, we observed that all aligners, even L-GRAAL that interchanges the input networks, obtained less than 10% of the edges matched in the target network.

**Table 2:** Edge Correctness scores obtained by the considered aligners.

| Net1 | Net2 | EC AligNet | EC HubAlign | EC L-GRAAL | EC PINALOG | EC SPINAL |
| --- | --- | --- | --- | --- | --- | --- |
| mus | cel | 0.58 | 0.81 | 0.79 | 0.34 | 0.01 |
| mus | sce | 0.65 | 0.97 | 0.68 | 0.56 | 0.05 |
| mus | dme | 0.65 | 0.88 | 0.70 | 0.30 | 0.03 |
| mus | hsa | 0.76 | 0.95 | 0.77 | 0.62 | 0.24 |
| cel | sce | 0.24 | 0.83 | 0.38 | 0.30 | 0.06 |
| cel | dme | 0.31 | 0.68 | 0.53 | 0.18 | 0.01 |
| cel | hsa | 0.31 | 0.77 | 0.43 | 0.23 | 0.01 |
| sce | dme | 0.03 | 0.01 | 0.08 | 0.19 | 0.03 |
| sce | hsa | 0.04 | 0.03 | 0.13 | 0.19 | 0.04 |
| dme | hsa | 0.13 | 0.37 | 0.31 | 0.13 | 0.01 |

**Table 3:** Functional Coherence scores obtained by the considered aligners.

| Net1 | Net2 | $FC_{max}$ | AligNet | HubAlign | L-GRAAL | PINALOG | SPINAL |
| --- | --- | --- | --- | --- | --- | --- | --- |
| | | | $FC - FC_{rel}$ | $FC - FC_{rel}$ | $FC - FC_{rel}$ | $FC - FC_{rel}$ | $FC - FC_{rel}$ |
| mus | cel | 0.21 | 0.06 – 0.26 | 0.04 – 0.21 | 0.03 – 0.17 | 0.10 – 0.50 | 0.12 – 0.60 |
| mus | sce | 0.24 | 0.08 – 0.33 | 0.07 – 0.30 | 0.04 – 0.18 | 0.12 – 0.50 | 0.15 – 0.64 |
| mus | dme | 0.19 | 0.05 – 0.24 | 0.03 – 0.16 | 0.03 – 0.13 | 0.07 – 0.42 | 0.06 – 0.33 |
| mus | hsa | 0.54 | 0.23 – 0.42 | 0.26 – 0.49 | 0.10 – 0.18 | 0.48 – 0.90 | 0.10 – 0.20 |
| cel | sce | 0.20 | 0.06 – 0.33 | 0.03 – 0.14 | 0.04 – 0.21 | 0.13 – 0.70 | 0.19 – 0.99 |
| cel | dme | 0.23 | 0.04 – 0.18 | 0.02 – 0.07 | 0.02 – 0.09 | 0.09 – 0.42 | 0.09 – 0.42 |
| cel | hsa | 0.24 | 0.04 – 0.17 | 0.02 – 0.08 | 0.03 – 0.13 | 0.08 – 0.35 | 0.08 – 0.36 |
| sce | dme | 0.24 | 0.05 – 0.19 | 0.07 – 0.30 | 0.02 – 0.07 | 0.07 – 0.31 | 0.10 – 0.43 |
| sce | hsa | 0.26 | 0.06 – 0.24 | 0.08 – 0.30 | 0.02 – 0.07 | 0.09 – 0.29 | 0.11 – 0.45 |
| dme | hsa | 0.20 | 0.04 – 0.18 | 0.02 – 0.11 | 0.02 – 0.08 | 0.09 – 0.41 | 0.08 – 0.43 |

Therefore, we can conclude that the analysis of the obtained results reveals that the EC score is only a measure of topological relation between the input networks. When the source network is smaller and has much less edges than the target network, then a high EC score should be expected. However, when the source network is similar to the target network or when it has a higher number of edges, then a very low EC score should be expected. Overall, it clearly implies that another definition of alignment correctness must be considered.

As far as the functional coherency goes, we report the obtained results in Table 3. We can observe there that all the aligners obtained a very low score. However, we cannot conclude that all aligners have a low biological coherence, because it is not clear if the low value is due to the alignment itself or to the measure of biological coherence. Therefore, we tried to obtain the maximum value of the FC score that can be expected for every pair of networks. To obtain this value we performed the following test: for every pair of input networks, we considered a complete bipartite graph where the nodes are the proteins in the two input networks, the edges are all protein pairs consisting on a protein in the source network and a protein in the target network, and the weight of each edge is the FC score of the corresponding pair of proteins. Then, we obtain the maximum FC score, *FC_max_*, as the FC score of the solution to the maximum weighted bipartite matching problem. Hence, for every pair of networks, we can compare the FC score obtained for each aligner with the maximum score. We define the *relative biological coherence* as the ratio between the FC score and the *FC_max_*. That is, 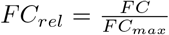. Now, if we look at the results presented in Table 3, we can observe that the values of *FC_max_* range from 0.19 to 0.26 and only in the alignment between *M. musculus* and *H. sapiens* we obtain that *FC_max_* = 0.54, which means that, in average in most of the alignments, the best alignment from the functional coherence point of view, maps correctly only 20% of the proteins. Considering now the results obtained by the different aligners, we can observe that the order from the highest to the lowest scores is almost the opposite to the order obtained when considering the EC scores. That is, the best scores are achieved by SPINAL, followed by PINALOG, AligNet, HubAlign and L-GRAAL. However, the results obtained even with SPINAL is that only 10% of the proteins is matched correctly, considering the GO term measure of biological coherence. In addition, the best possible alignment would be able to align correctly only 20% of the proteins. We discard the hypothesis that the GO term measure is low due to the lack of GO terms, since, as we show in Table 4, for every pair of networks, 90% of protein pairs have their GO terms annotated, and there is no correlation between the number of annotated GO terms and FC scores. Therefore, since the FC scores are not conclusive of meaningful biological alignments, we performed an additional test explained below.

**Table 4:** This table shows the relation between available GO pairs for every pair of networks.

| Net1 | Net2 | $FC_{max}$ | NotAvailableGOpairs | AvailableGOpairs | Percentage |
| --- | --- | --- | --- | --- | --- |
| mus | cel | 0.21 | 1 | 598 | 99.83 |
| mus | sce | 0.24 | 0 | 599 | 100.00 |
| mus | dme | 0.19 | 2 | 597 | 99.67 |
| mus | hsa | 0.54 | 1 | 598 | 99.83 |
| cel | sce | 0.20 | 20 | 2954 | 99.33 |
| cel | dme | 0.23 | 256 | 2718 | 91.39 |
| cel | hsa | 0.24 | 102 | 2872 | 96.57 |
| sce | dme | 0.24 | 45 | 5478 | 99.19 |
| sce | hsa | 0.26 | 14 | 5509 | 99.75 |
| dme | hsa | 0.20 | 0.19 | 7126 | 96.47 |

#### Qualitative analysis

As stated in [8], the evaluation of any aligner should be done considering also their quality and not only the quantity, that is, the accuracy of the alignment and not only the number of preserved edges or GO terms. The accuracy of any aligner is easily tested when there is a gold standard to compare to. With this idea in mind, we have considered the following test to study the quality of our alignments in contrast with the quality of the alignments obtained by PINALOG, HubAlign and L-GRAAL. Since the results obtained by SPINAL in the previous test were not convincing, we do not consider it in the next test.

#### Protein complex prediction

In order to test the behavior of AligNet in the alignment of protein complexes, we also performed the protein complex prediction test reported in [30] using PINALOG. Following the procedure explained therein, we considered the database MIPS CORUM [33] for the human protein complexes and, as a gold standard for the yeast complexes, we considered the information available in [11]. In addition, we considered the functional information available in MIPS CORUM for the human complexes and in MIPS FunCat [34] for the yeast complexes. Then, we considered the overlapping score of complexes introduced in [30], and we defined a functional coherence value for the alignment of protein complexes, as explained below.

Let *G* = (*V*, *E*) and *G*′ = (*V*′,*E*′) be two PPIN such that |*V*| ⩽ |*V*′|, let *μ* : *G* → *G*′ be a mapping, and let *c* ⊆ *V* and *c*′ ⊆ *V*′ be two protein complexes in *G* and *G*′, respectively. The *overlapping score of c and c*′ is defined as

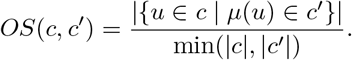

To define a functional coherence value for the protein complex alignment, every protein complex *c* in *G* is first mapped to a protein complex *c*′ in *G*′ provided that *OS*(*c, c*′) lies over a threshold, fixed at 0.2 as done in [30]. Next, for every complex c in G we consider the complex *c*′ in *G*′ such that *OS*(*c, c*′) is maximum.

As it was the case with the edge correctness ratio, an overlapping score of pairs of complexes of 100% need not be evidence of a correct network alignment, because every protein complex is supposed to develop several biological functions, and the alignment may establish a correspondence between two complexes that are completely unrelated from the point of view of their function.

The main point here is that the aim of the alignment should be clearly stated. If it only aims at matching similar topological substructures of the networks, in order to detect those substructures that appear in both networks, then maximizing the sum of the overlapping score of pairs of complexes may be a suitable goal. However, if the alignment searches for pairs of proteins that share biological functions, then only those complexes with a common function should be matched. Since the main application of PPIN alignment is to infer biological functions of proteins and protein complexes, it is very important that the alignment does not match biologically unrelated complexes. Therefore, we define the complex functional coherence of an alignment between PPIN as follows. First, a pair of two complexes, one in each network, is said to be *coherent* if they share some biological function; otherwise, the pair is incoherent. Then, the *complex functional coherence value* (CFC) of the alignment is defined by the complex alignment precision, that is, the ratio of complexes that are aligned correctly with respect to the aligned complexes. If we denote by CP the number of coherent pairs and by NCP the number of incoherent pairs, then 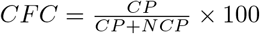.

In Table 5 we show the results obtained by all the aligners. We can observe there that AligNet does not align 1269 complexes and produces 377 incoherent pairs and 128 coherent pairs. HubAlign does not align 1154 complexes and produces 589 incoherent pairs and 31 coherent pairs. PINALOG does not align 945 complexes and produces 626 incoherent pairs and 203 coherent pairs. Finally, L-GRAAL does not align 996 complexes and produces 741 incoherent pairs and 31 coherent pairs. Thus, the CFC value obtained in the alignment produced by AligNet is 25.34, which means that AligNet aligns correctly 25% of the assigned complexes. Very close to AligNet is the CFC value in the alignment produced by PINALOG, which is 24.48. However, the CFC values obtained in the alignments produced by L-GRAAL and HubAlign are lower. They are 4.75 and 5 respectively, which means that they align correctly 5% of the matched complexes.

**Table 5:** This table shows the number of complexes that are not assigned/assigned correctly and assigned incorrectly.

|  | AligNet | HubAlign | PINALOG | L-GRAAL |
| --- | --- | --- | --- | --- |
| NotAssigned | 1269 | 1154 | 945 | 996 |
| NotCoherent | 377.00 | 589.00 | 626.00 | 741 |
| Coherent | 128 | 31 | 203 | 37 |
| CFC | 25.34 | 5 | 24.48 | 4.75 |

In order to analyze the results obtained by AligNet in contrast to the others aligners, we also considered the following metrics: to contrast AligNet versus another aligner, for instance HubAlign, we first count the number of complexes that are not aligned either by AligNet nor by HubAlign. Then, for those complexes that are aligned by AligNet and are not aligned by HubAlign, we count the number of coherent and incoherent pairs. Conversely, for those complexes that are not aligned by AligNet and are aligned by HubAlign, we count the number of coherent and incoherent pairs. We show these results in Table 6. We can see there that there are 891 complexes that are not aligned neither by AligNet nor by HubAlign. This means that 77% of the complexes that are not aligned by AligNet are also not aligned by HubAlign. With respect to the remaining 263 complexes that are aligned by AligNet but not by HubAlign, 88 are correctly aligned (coherent pairs) and 175 are incorrectly aligned. On the other hand, there are 378 complexes that are aligned by HubAlign but not by AligNet, from those, 21 are correctly aligned and 357 produced incoherent pairs. Therefore, we can conclude that HubAlign assigned more complexes than AligNet, but, when we look in detail to the complexes alignment, AligNet was able to align correctly 88 complexes that HubAlign did not align. HubAlign aligned correctly only 21 complexes that AligNet did not align. In addition, HubAlign aligned incorrectly 357 complexes that AligNet did not align and AligNet aligned incorrectly 175 complexes that HugAlign did not align. This means that AligNet achieves a higher precision in complex alignment than HubAlign.

**Table 6:** This table shows how AligNet assigned the complexes that are not assigned by HubAlign and conversely.

|  | AligNet vs HubAlign | Ratio | HubAlign vs AligNet | Ratio |
| --- | --- | --- | --- | --- |
| Not Assigned | 891 | 77.21 | 891 | 70.21 |
| Not Coherent | 175 | 15.16 | 357 | 28.13 |
| Coherent | 88 | 7.63 | 21 | 1.65 |

Concerning the results shown in Table 7, when we contrast AligNet with L-GRAAL, we obtain a similar result to the contrast of AligNet with HubAlign. Again, AligNet obtains a higher precision than L-GRAAL since it aligned correctly 66 complexes and incorrectly 167, while L-GRAAL aligned correctly 29 complexes and incorrectly 477.

**Table 7:** This table shows how AlignNet assigned the complexes that are not assigned by HubAlign and conversely.

|  | AligNet vs L-GRAAL | Ratio | L-GRAAL vs AligNet | Ratio |
| --- | --- | --- | --- | --- |
| Not Assigned | 763 | 76.61 | 763 | 60.13 |
| Not Coherent | 167 | 16.77 | 477 | 37.59 |
| Coherent | 66 | 6.63 | 29 | 2.29 |

However, the results obtained when we contrast AligNet with PINALOG show that, indeed, they obtain a similar precision in complex alignment (See Table 8). We obtain that AligNet aligned correctly 25 complexes and incorrectly 105 while PINALOG aligned correctly 79 complexes and incorrectly 375. Thus, the ratio of coherent pairs over assigned complexes is similar, 0.19 for AligNet and 0.17 for PINALOG. Therefore, the only differences that can be observe between these aligners is that PINALOG aligns more complexes than AligNet. If it aligns more complexes, and its precision is slightly the same as the precision of AligNet, then PINALOG has more incorrect alignments than AligNet and also more correct alignments. In this sense, AligNet is a more conservative aligner than PINALOG, although its precision is slightly higher than the precision of PINALOG.

**Table 8:** This table shows how AligNet assigned the complexes that are not assigned by PINALOG and conversely.

|  | AlignNet vs PINALOG | Ratio | PINALOG vs AlignNet | Ratio |
| --- | --- | --- | --- | --- |
| Not Assigned | 815.00 | 86.24 | 815.00 | 64.22 |
| Not Coherent | 105 | 11.11 | 375 | 29.55 |
| Coherent | 25 | 2.65 | 79 | 6.23 |

We present in Figure 8 a visualization of the obtained results when we contrasted AligNet with the other aligners. We present there the ratio of unaligned complexes, correctly aligned complexes (coherent pairs) and incorrectly aligned complexes (incoherent pairs). We can observe that HubAlign versus AligNet (second bar from the left) as well as L-GRAAL versus AligNet (first bar from the right) obtain a higher proportion of incoherent pairs and a lower proportion of coherent pairs. In contrast, AligNet versus PINALOG and PINALOG verus AligNet (the two bars in the center) obtain a similar proportion of correctly and incorrectly aligned pairs.

**Figure 8:**
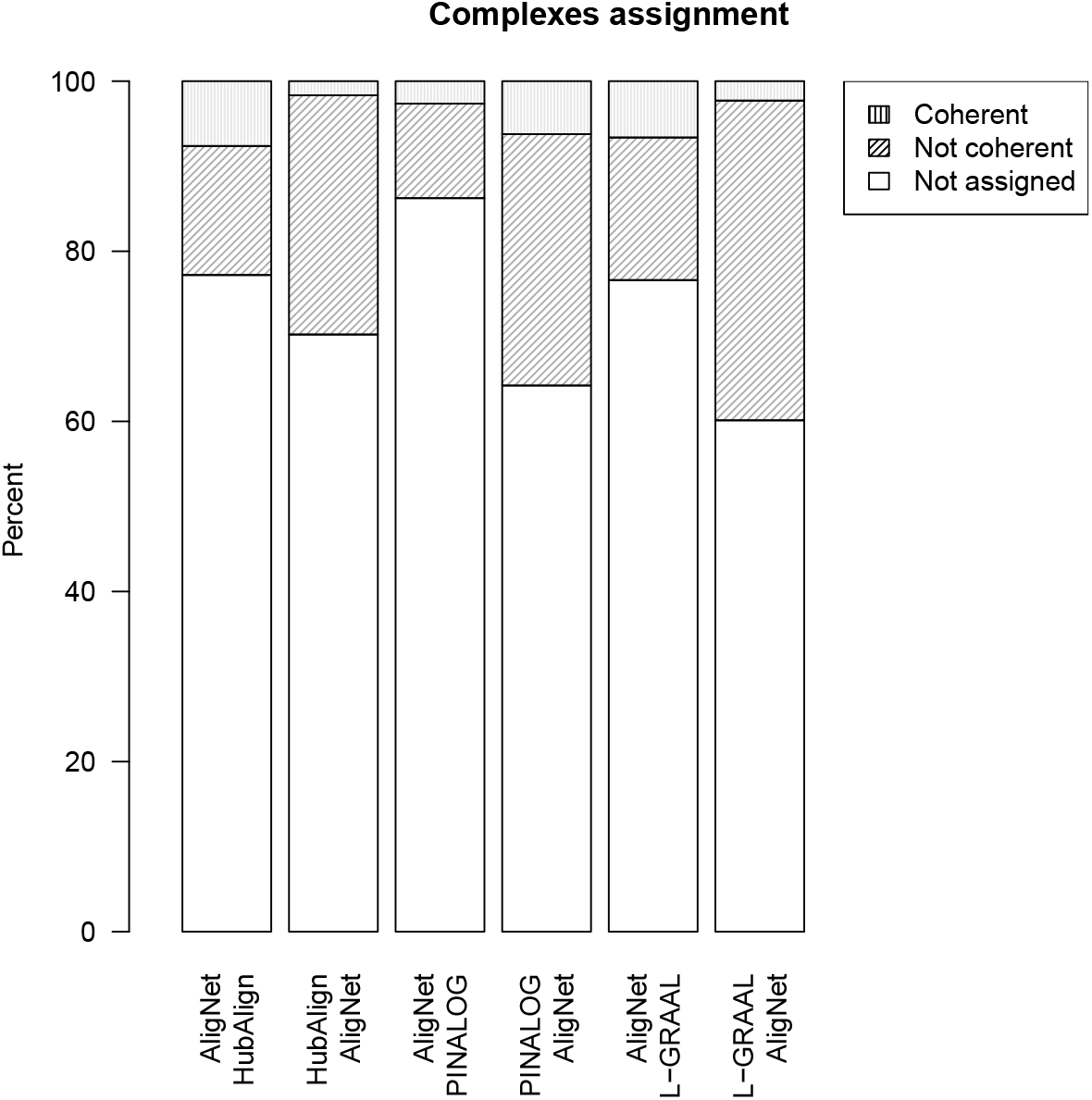
This figure shows the results in the protein complexes test obtained by AligNet in contrast to the others aligners, when we consider the alignment of complexes between S. cerevisiae and H. sapiens. We show the proportion obtained by AligNet of coherent, not coherent and not assigned complexes when the other aligners do not assign complexes, and conversely. Thus, the first bar shows the proportion between coherent, not coherent and not assigned complexes by AligNet when HubAlign does not assigned. Conversely, the second bar shows the proportion between coherent, not coherent and not assigned complexes by HubAlign when AligNet does not assigned.

As a result of the comparison between the aligners, we obtain again, as it was the case in [8] and [24], that the agreement of the alignments obtained by different aligners is vey low. The majority of the global aligners achieve a high node coverage, meaning that the average of assigned nodes in the source network is high, but all of them obtain a very low biological coherence value. With respect to the topological coherence value, some aligners are able to obtain a high score but it is always associated with a biological coherence score. Overall, we can conclude that AligNet is the aligner that obtains a better balance between topological coherency (it preserves 60% of the edges) and functional coherency (relative function coherency values between 20% and 40% and the highest complex functional coherency score, 25.34), followed by PINALOG, which obtains similar functional coherency scores than AligNet and a bit lower topological coherency scores. However, HubAlign and L-GRAAL obtain high topological coherency scores but low functional coherency values. On the other hand, SPINAL surprisingly obtains a very low topological coherency value. Thus, if the purpose of the alignment is to align correctly, in the biological function sense, we propose to adopt AligNet since it is the most precise. However, if the purpose of the alignment is to find some topological similarity, then we propose to adopt HubAlign.

### Aligners analysis

In order to study the efficiency of the considered aligners, we take into account their running time and memory space needed to perform an alignment. We run our implementation of AligNet on a server with 4 processors at 2.6 GHz and 20 GB of RAM and we also run the latest implementation of NATALIE (downloaded from http://www.mi.fu-berlin.de/w/LiSA/Natalie), PINALOG (downloaded from http://www.sbg.bio.ic.ac.uk/~PINALOG/), SPINAL (downloaded from http://code.google.com/p/spinal/), HubAlign (downloaded from http://ttic.uchicago.edu/hashemifar/software/HubAlign.zip) and L-GRAAL (downloaded from http://bio-nets.doc.ic.ac.uk/L-GRAAL/).

As we already explained in the Background section, one of the weak points of PPIN aligners is either their running time or the memory space they use. Indeed, although NATALIE was suggested as a good aligner, it could not even align the two smallest networks, *C. elegans* and *D. melanogaster*, on a computer with 64 GB of RAM. With respect to PINALOG, SPINAL, HubAlign and L-GRAAL, we were able to complete all the alignments and we show their running times in Table ??. In order to visualize their running times, we also show the running times of every finished computation for each aligner in Figure 9. We can observe there that SPINAL is, with a big difference, the slowest one to compute the alignments between *H. sapiens* and *S. cerevisiae*, and also between *D. melanogaster* and *S. cerevisiae*. In addition, PINALOG is the slowest one, also with a big difference, to compute the alignment between *C. elegans* and *H. sapiens*, as well as the alignment between *H. sapiens* and *M. musculus*. We can also observe that AligNet is considerably faster than PINALOG and SPINAL, with a running time of less than a thousand seconds in most of the alignments. Only in one computation, the alignment between *D. melanogaster* and *H. sapiens*, AligNet is slower than PINALOG and SPINAL with a difference of less than two thousand seconds. However, it is difficult to see the running times in some alignments because SPINAL needed more than 20,000 seconds for the alignment between *S. cerevisiae* and *H. sapiens*. Thus, in order to visualize the results in the cases where the aligners consumed less than 3,500 seconds, we decided to remove SPINAL and PINALOG and in Figure 10 we show agin the results considering only AligNet, HubAlign and L-GRAAL. We can observe there that L-GRAAL is the aligner that consumed more time in most of the computations. Concerning HubAlign and AligNet, HugAlign is faster except in the alignments between *C. elegans* and *S. cerevisiae* and also *S. Cerevisiae* and *D. melanogaster*.

**Figure 9:**
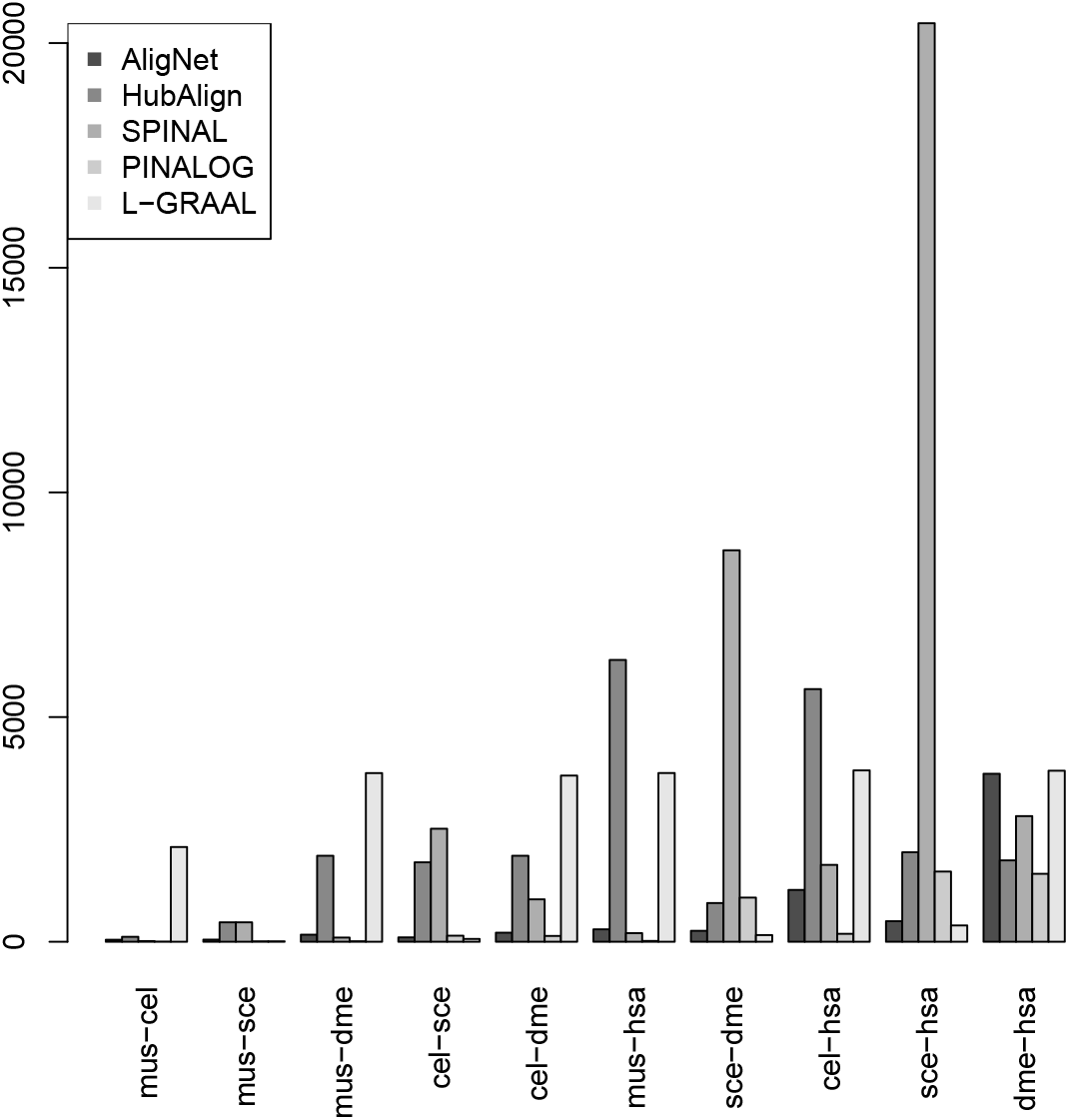
This figure shows the running times (in seconds) we obtained when we performed all the alignments for every pair of the considered networks. In the figure we present the results obtained with the aligners AligNet, PINALOG, SPINAL, HubAlign and L-GRAAL.

**Figure 10:**
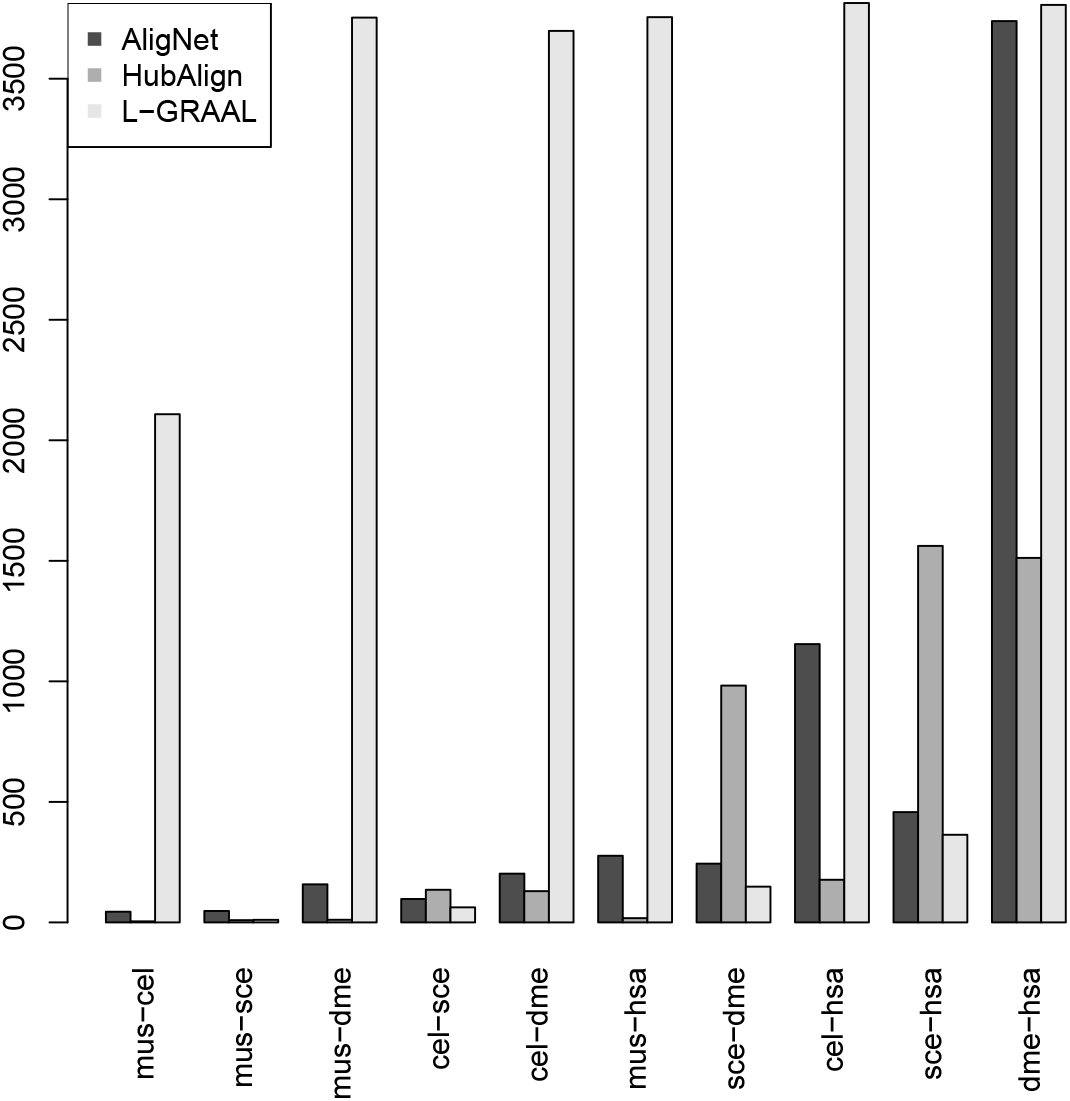
This figure shows the same information presented in Figure 9 considering only the aligners AligNet, HubAlign and L-GRAAL.

Furthermore, we show in Figure 11 the relation between network size and running time for all of the computations with each of the aligners. The size of a network pair is the sum of their nodes. Thus, the network pairs in the diagrams are positioned in increasing order. A perfect aligner, from the efficiency point of view, should present a linear relation between the size of the network pair and the consumed time. From top left to bottom right, we show the results for the aligners AligNet, HubAlign, SPINAL, PINALOG, and L-GRAAL. We can observe that HubAlign and AligNet present a clear relation between computation time and size of the input networks. However, this is not the case of PINALOG, SPINAL and L-GRAAL. It should be noticed here, that L-GRAAL has a step parameter which may force to stop the computation.

**Figure 11:**
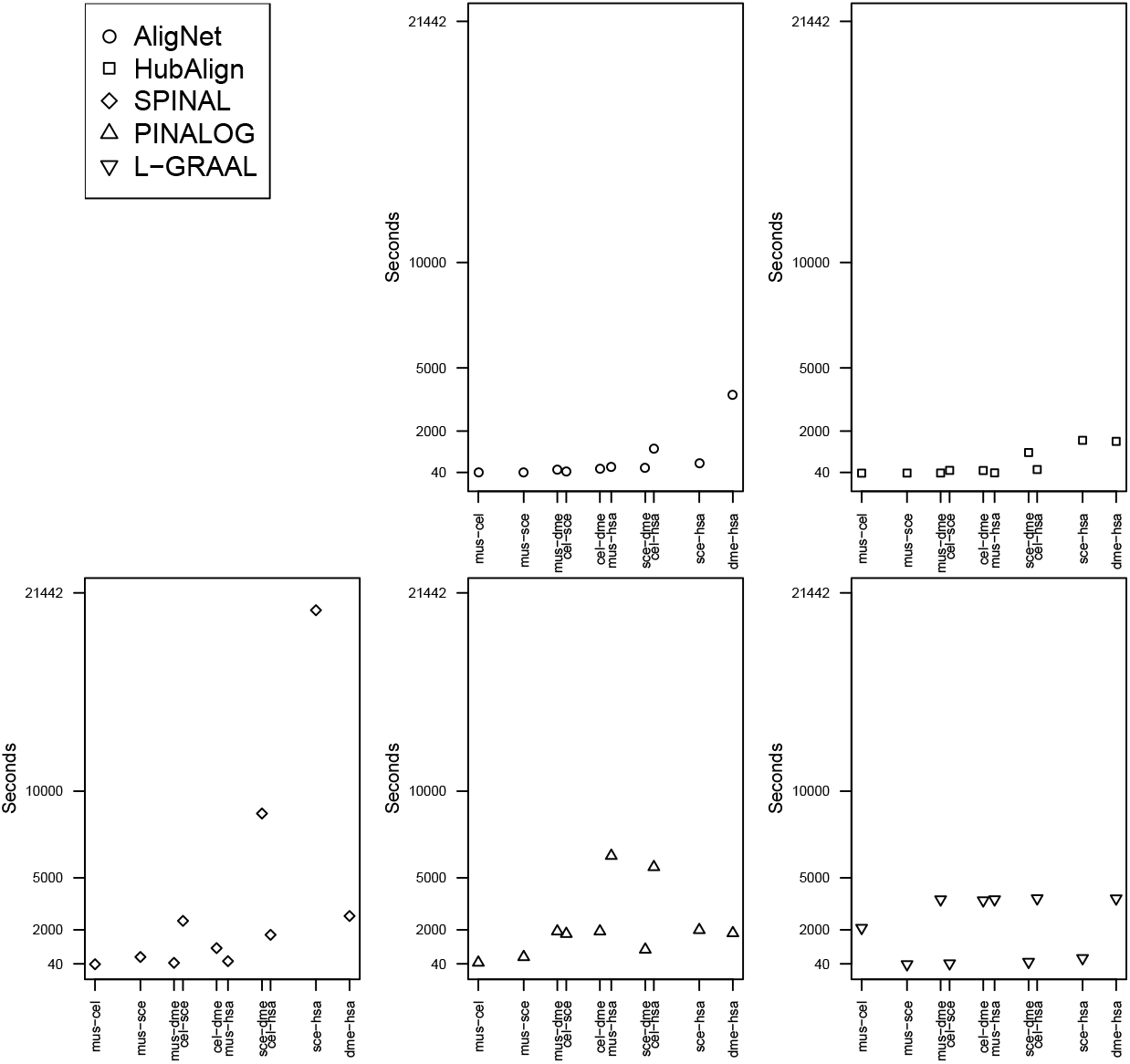
This figure shows the relation between the time needed to obtain every alignment and the size of the input networks with the aligners AligNet, PINALOG, SPINAL, HubAlign and L-GRAAL.

## Conclusions

In this paper we present AligNet, a new method and software tool for the pairwise global alignment of PPIN aimed to produce biologically meaningful alignments by achieving a good balance between structural matching and protein function conservation. AligNet is a parameter-free algorithm that, given two PPIN, produces a consistent alignment from the smaller network, in terms of number of nodes, to the larger network. In order to assess the correctness of our AligNet aligner, we have evaluated the quality of the alignments obtained with AligNet and with the best aligners established in [8, 24], namely: PINALOG, SPINAL, HubAlign, and L-GRAAL. The obtained results show that, indeed, AligNet produces biologically more meaningful alignments than state-of-the-art methods and tools, by achieving a better balance between structural matching and protein function conservation.

We have used both the edge correctness (EC) and the functional coherence (FC) metrics. The results obtained and presented in the Results and Discussion section of this paper, reveal that HubAlign and L-GRAAL obtained the best EC scores when the source input network has considerably less number of edges than the target network, preserving 80% of the edges, while AligNet preserved 60% of the edges. However, all aligners obtained very low EC scores, with less than 10% of the edges preserved, when the input PPIN have similar size.

Concerning the efficiency of the considered aligners from the computational point of view, HubAlign and AligNet obtained the best running time. In addition, running time increases with both aligners with the increase in the size of the input networks, unlike the other aligners, which are slower than HubAlign and AligNet and have a variable running time that is not related to the size of the input networks.

## Acknowledgements

We thank Gabriel Riera for the technical support.

## References

[1] Ahmet E Aladag and Cesim Erten. SPINAL: scalable protein interaction network alignment. Bioinformatics, 29(7):917–924, 2013.

[2] Ahmet E. Aladağ and Cesim Erten. SPINAL: Scalable protein interaction network alignment. Bioinformatics, 29(7):917–924, 2013.

[3] Michael Ashburner et al. Gene Ontology: tool for the unification of biology. Nat. Genet., 25:25–29, 2000.

[4] Ralf Borndörfer and Olga Heismann. The hypergraph assignment problem. Discrete Optim., 15:15–25, 2015.

[5] Christiam Camacho, George Coulouris, Vahram Avagyan, Ning Ma, Jason Papadopoulos, Kevin Bealer, and Thomas L. Madden. BLAST+: architecture and applications. BMC Bioinformatics, 10(1):1, 2009.

[6] L. Chindelevitch, CY. Ma, CS. Liao, and B. Berger. Optimizing a global alignment of protein interaction networks. Bioinformatics, 29(21):2765–73, 2013.

[7] C. Clark and J. Kalita. A multiobjective memetic algorithm for ppi network alignment. Bioinformatics, 31(12):1988–98, 2015.

[8] Connor Clark and Jugal Kalita. A comparison of algorithms for the pairwise alignment of biological networks. Bioinformatics, 30(16):2351–2359, 2014.

[9] Ahed Elmsallati, Connor Clark, and Jugal Kalita. Global alignment of protein-protein interaction networks: A survey. IEEE/ACM Transactions on Computational Biology and Bioinformatics, 13(4):689–705, 2016.

[10] Jason Flannick, Antal Novak, Balaji S Srinivasan, Harley H. McAdams, and Serafim Batzoglou. Graemlin: general and robust alignment of multiple large interaction networks. Genome Res., 16(9):1169–81, 2006.

[11] Anne-Claude Gavin, Patrick Aloy, Paola Grandi, Roland Krause, Markus Boesche, Martina Marzioch, Christina Rau, Lars Juhl Jensen, Sonja Bastuck, Birgit Dümpelfeld, et al. Proteome survey reveals modularity of the yeast cell machinery. Nature, 440(7084):631–636, 2006.

[12] Pietro Hiram Guzzi and Tijana Milenković. Survey of local and global biological network alignment: the need to reconcile the two sides of the same coin. Briefings in bioinformatics, page bbw132, 2017.

[13] S. Hashemifar, J. Ma, H. Naveed, S. Canzar, and J. Xu. Modulealign: module-based global alignment of protein-protein interaction networks. Bioinformatics, 32(17):658–64, 2016.

[14] S. Hashemifar and J. Xu. HubAlign: an accurate and efficient method for global alignment of protein-protein interaction networks. Bioinformatics, 30(17):i438–i444, 2014.

[15] Nazlin K Howell. Protein-protein interactions. In Biochemistry of food proteins, pages 35–74. Springer, 1992.

[16] Brian P Kelley, Bingbing Yuan, Fran Lewitter, Roded Sharan, Brent R Stockwell, and Trey Ideker. PathBLAST: a tool for alignment of protein interaction networks. Nucleic Acids Res., 32(Web Server issue):W83–88, July 2004.

[17] Ozlem Keskin, Nurcan Tuncbag, and Attila Gursoy. Predicting protein-protein interactions from the molecular to the proteome level. Chemical Reviews, 116(8):4884–4909, 2016.

[18] Hiroaki Kitano. Systems biology: a brief overview. Science, 295(5560):1662–1664, 2002.

[19] Gunnar W. Klau. A new graph-based method for pairwise global network alignment. BMC Bioinformatics, 10(Suppl. 1):S59, 2009.

[20] Mehmet Koyutürk, Yohan Kim, Umut Topkara, Shankar Subramaniam, Wojciech Szpankowski, and Ananth Grama. Pairwise alignment of protein interaction networks. J. Comput. Biol., 13(2):182–199, March 2006.

[21] HW Kuhn. The Hungarian method for the assignment problem. Naval Research Logistics, 52(1):7–21, 2005.

[22] Zhenping Li, Yong Wang, Shihua Zhang, Xiang-Sun Zhang, and Luonan Chen. Alignment of protein interaction networks by integer quadratic programming. In Proc. 28th Annual International Conference of the IEEE Engineering in Medicine and Biology Society, pages 5527–5530, 2006.

[23] Zhi Liang, Meng Xu, Maikun Teng, and Liwen Niu. NetAlign: a web-based tool for comparison of protein interaction networks. Bioinformatics, 22(17):2175–2177, September 2006.

[24] N. Malod-Dognin, K. Ban, and N. Pržulj. Unified alignment of protein-protein interaction networks. Scientific Reports, 7(953), 2017.

[25] N. Malod-Dognin and N. Przulj. L-graal: Lagrangian graphlet-based network aligner. Bioinformatics, 31(13):2182–9, 2015.

[26] Manikandan Narayanan and Richard M. Karp. Comparing protein interaction networks via a graph match-and-split algorithm. J. Comput. Biol., 14(7):892–907, September 2007.

[27] Behnam Neyshabur, Ahmadreza Khadem, Somaye Hashemifar, and Seyed Shahriar Arab. NETAL: a new graph-based method for global alignment of protein-protein interaction networks. Bioinformatics, 29(13):1654–1662, 2013.

[28] Daniel Park, Rohit Singh, Michael Baym, Chung-Shou Liao, and Bonnie Berger. IsoBase: a database of functionally related proteins across PPI networks. Nucleic Acids Res., 39(suppl 1):D295–D300, 2011.

[29] Rob Patro and Carl Kingsford. Global network alignment using multiscale spectral signatures. Bioinformatics, 28(23):3105–3114, 2012.

[30] Hang T. T. Phan and Michael J. E. Sternberg. PINALOG: A novel approach to align protein interaction networks—implications for complex detection and function prediction. Bioinformatics, 28(9):1239–1245, 2012.

[31] R Core Team. R: A Language and Environment for Statistical Computing. R Foundation for Statistical Computing, Vienna, Austria, 2015.

[32] V. Srinivasa Rao, K. Srinivas, G. N. Sujini, and G. N. Sunand Kumar. Protein-protein interaction detection: Methods and analysis. Int. J. Proteomics, 2014:147648, 2014.

[33] Andreas Ruepp, Barbara Brauner, Irmtraud Dunger-Kaltenbach, Goar Frishman, Corinna Montrone, Michael Stransky, Brigitte Waegele, Thorsten Schmidt, Octave Noubibou Doudieu, Volker Stümpflen, and H. Werner Mewes. CORUM: the comprehensive resource of mammalian protein complexes. Nucleic Acids Res., 36(suppl 1):D646–D650, 2008.

[34] Andreas Ruepp, Alfred Zollner, Dieter Maier, Kaj Albermann, Jean Hani, Martin Mokrejs, Igor Tetko, Ulrich Güldener, Gertrud Mannhaupt, Martin Munsterkötter, and H. Werner Mewes. The FunCat, a functional annotation scheme for systematic classification of proteins from whole genomes. Nucleic Acids Res., 32(18):5539–5545, 2004.

[35] Andreas Schmidt, Ignasi Forne, and Axel Imhof. Bioinformatic analysis of proteomics data. BMC Syst. Biol., 8(Suppl 2):S3, 2014.

[36] Rohit Singh, Jinbo Xu, and Bonnie Berger. Global alignment of multiple protein interaction networks with application to functional orthology detection. Proceedings of the National Academy of Sciences of the United States of America, 105(35):12763–12768, 2008.

[37] V. Vijayan, V. Saraph, and T. Milenković. Magna++: Maximizing accuracy in global network alignment via both node and edge conservation. Bioinformatics, 31(14):2409–11, 2015.

